# Feeding frequency sets the rhythm of asexual reproduction in the sea anemone *Cylista elegans*

**DOI:** 10.64898/2026.07.13.738267

**Authors:** C. D. Wells, L. G. Harris

## Abstract

Many sea anemones reproduce asexually by pedal laceration, shedding fragments of the pedal disk that regenerate into polyps. How feeding affects the amount of this reproduction differs among species, but whether feeding also sets its timing has rarely been examined. We fed the sea anemone *Cylista elegans* (formerly *Sagartia elegans*) daily, every second day, every fourth day, or not at all, measured growth, laceration, and survival over 35 days, then reassigned anemones to new schedules to test whether laceration follows the current or previous schedule. Growth rose with feeding but saturated, animals fed daily and every second day growing at similar rates. Total laceration depended on whether an anemone was fed rather than how often. Every fed schedule produced more lacerates than starvation. The timing, by contrast, tracked the schedule closely. Laceration was suppressed for about a day after each meal and recovered before the next, so anemones fed every second or every fourth day lacerated on matching two- and four-day rhythms, while daily-fed animals were arrhythmic. When moved to a new schedule, the period shifted to match it, not the old one. Only starvation caused death. Asexual reproduction in *C. elegans* is therefore bound closely to feeding, which fuels laceration yet suppresses it during digestion, confining it between meals. This coupling distinguishes *C. elegans* from anemones with symbiotic microalgae, where starvation rather than feeding drives reproduction, and links it to the feeding-driven clonal proliferation of the invasive sea anemone *Diadumene lineata*.

## 1. Introduction

Many animals reproduce not only by sex but by cloning themselves, splitting, budding, or shedding pieces of the body that regrow into whole new individuals. This capacity is well dispersed across the animal tree, in sponges, cnidarians, flatworms, annelids, bryozoans, echinoderms, and colonial ascidians, and is richest in soft-bodied groups that carry no rigid skeleton to break (Blackstone and Jasker, 2003; Fautin, 2002; Jackson, 1986). In such animals, which often grow without a fixed adult size, the line between growing and reproducing is thin: tissue, shed as a new clonemate, is energy the parent allocates to neither its own growth nor its gametes, and how an animal divides its intake among growing, cloning, and sexually reproducing, and when, is set by the food and conditions around it (Sebens, 1979, 1987). Sea anemones make this trade through pedal laceration, fission, or budding. In pedal laceration, small pieces of the pedal disk tear away as the animal moves and regenerate into new polyps (Cary, 1911; Stephenson, 1929; van der Burg and Prentis, 2021); laceration begins where part of the disk stays fixed while the column moves away, stretching the tissue until it ruptures and leaves a fragment behind to regenerate (Cary, 1911; Geller et al., 2005). How often an anemone lacerates is not fixed but tracks the environment: temperature, immersion, water flow, food supply, substrate, and crowding each change the rate at which a clone divides (Al-Shaer et al., 2023; Anthony and Svane, 1994; Geller et al., 2005; Johnson and Shick, 1977; Sebens, 1980), letting a single genotype adjust its output of clonemates to local conditions.

Among these factors, food has drawn the most attention, and its effect on asexual reproduction differs sharply among species. In the sea anemones *Exaiptasia diaphana* (formerly *Aiptasia*) and *Anthopleura elegantissima* that host endosymbiotic microalgae, frequent feeding suppresses asexual reproduction: well-fed animals grow but seldom divide, whereas starvation has been reported to trigger asexual reproduction (Clayton, 1985; Sebens, 1980). The response is not uniform even among these species, however: feeding has been reported to increase fission in *A. elegantissima* (Tsuchida and Potts, 1994), contrary to findings by Sebens (1980). Nutritional state also sets the energy reserves an animal carries into each new lacerate (Clayton, 1985; White et al., 2026). The opposite pattern holds in species that divide as part of growth rather than in response to scarcity. Fission in *Diadumene lineata* (formerly *Haliplanella luciae*) and clone formation in the moon jelly *Aurelia aurita* both accelerate as feeding becomes more frequent (Keen and Gong, 1989; Minasian, 1976). How feeding maps onto asexual reproduction is therefore species-specific and, for most anemones, still poorly resolved.

Almost all of this work concerns how much an anemone reproduces, not when. Yet feeding in the field is episodic, and where laceration depends on energy balance it might be timed to the interval between meals. The few studies to examine this timing found it to be species-specific as well. *Aiptasiogeton hyalinus* lacerates on the days farthest from its last feeding, whereas *E. diaphana* lacerates at random between meals (Clayton, 1985; Smith and Lenhoff, 1976). Anemones maintain endogenous circadian rhythms of activity and metabolism, which are entrained by external cues and persist when those cues are removed (Hendricks et al., 2012; Maas et al., 2016; Oren et al., 2015). Whether the rhythm of asexual reproduction likewise entrains to the rhythm of feeding, and whether such a rhythm is fixed by past feeding or reset by current feeding, has not been tested.

*Cylista elegans* (formerly *Sagartia elegans*) is well suited to these questions: it reproduces asexually only by pedal laceration and lacerates readily in the laboratory. It is an acontiate species, native to the European coast, that forms localized clones spreading over tens of meters (Grebelnyi and Kovtun, 2013; Shaw, 1991; Stephenson, 1929). An introduced population persisted for roughly a decade at a single marina in Salem Harbor, Massachusetts, the only record of the species in the western Atlantic, before disappearing after the winter of 2010 to 2011 (Pederson et al., 2005; Wells and Harris, 2019). The causes of that disappearance were primarily thermal and are reported elsewhere (Wells, 2013; Wells and Harris, 2019); here we use animals from the same population.

We examine how feeding frequency affects the growth, asexual reproduction, and survival of *C. elegans*, and whether the timing of pedal laceration entrains to the feeding schedule. We fed anemones daily, every second day, every fourth day, or not at all, and then reassigned them to new schedules in a second trial to test whether any rhythm of reproduction follows the current feeding schedule or carries the history of the previous one. We hypothesized that more frequent feeding would increase growth and total reproduction and that pedal laceration would form a rhythm matching the feeding interval.

## 2. Materials and methods

We collected *C. elegans* in October 2010 from floating docks at Hawthorne Cove Marina, Salem, Massachusetts, USA (42°31′ N, 70°52′ W). Polyps were detached by hand from the shells of the blue mussel *Mytilus edulis* and held at 15 °C on a 12:12 h light:dark cycle in unfiltered seawater drawn from the University of New Hampshire Coastal Marine Laboratory, New Castle, New Hampshire, USA (43°4′ N, 70°42′ W). Animals were fed nauplii of the brine shrimp *Artemia franciscana*.

We tracked growth as pedal disk surface area (PDSA), a non-destructive measure that let each polyp remain in the experiment undisturbed. PDSA was calculated as the area of an ellipse, (π/4) · *D*_max_ · *D*_perp_, where *D*_max_ is the maximum diameter of the pedal disk and *D*_perp_ is the diameter perpendicular to it. Pedal disks in this species are frequently elongated rather than circular, their two diameters differing severalfold, so an ellipse approximates their area far better than a circle would. Calculated PDSA closely predicted disk area measured directly from calibrated photographs (weighted least-squares regression, R^2^ > 0.99, Wells, 2013), so the two diameters were sufficient to follow growth.

Two hundred polyps (maximum pedal-disk diameter 1.26 to 5.90 mm, mean 3.04 mm) were distributed evenly among forty 1.2-L containers, five polyps per container, each in 1.0 L of unfiltered seawater in a temperature-controlled room at 15 °C. After a seven-day starvation period that cleared residual food and set a common baseline, containers were assigned to one of four feeding schedules: fed daily, every second day, every fourth day, or left unfed, with ten containers per schedule. At each feeding, 1.0 mL of *A. franciscana* nauplii (approximately 24,000 nauplii mL^-1^) was added and the polyps were allowed to feed for 2 h, well beyond repletion. After feeding, water of all containers was replaced with unfiltered seawater. Pedal lacerates were counted and removed once each day, on feeding days immediately before that day’s feeding, so that each count records the lacerates produced over the preceding 24 h. PDSA was measured at the start of the experiment and every seven days through day 35, and mortality was recorded daily. These measurements constitute the first trial.

After 35 days, the thirty previously fed containers were each reassigned at random to daily, every second day, every fourth day, or unfed, to test whether the reproductive rhythm follows the current schedule or carries the previous one. Husbandry, lacerate counts, and PDSA measurements continued as in the first trial for a further 35 days. These measurements constitute the second trial.

All analyses were performed in R 4.6.0 (R Core Team, 2026); the data and code are archived at Zenodo (Wells and Harris, 2026). Throughout, the experimental unit was the container: the ten containers per schedule were the replicates, not the individual anemones within a container as they were not independent of one another. For each container, we measured the anemones’ growth as the slope of their PDSA against day and compared these growth rates among schedules by analysis of variance with Tukey’s post-hoc comparisons, and again with a linear mixed model that let each container have its own starting size (R package lme4 2.0.1, Bates et al., 2015; R package emmeans 2.0.3, Lenth and Piaskowski, 2026). We also repeated the growth comparison on proportional growth and with starting size as a covariate, to confirm that any similarity among schedules was not an artifact of the growth measure or of starting size. Growth in the second trial was analyzed against both the first-trial and second-trial schedules. Feeding efficiency was a container’s total growth divided by the number of feedings it received. We reconstructed each polyp’s death or censoring time from the daily survivor counts and summarized survival with the Kaplan-Meier estimator (R package survival 3.8.6, Therneau, 2026).

We modeled the total lacerates each container produced over 35 days with a negative-binomial model (R package glmmTMB 1.1.14, Brooks et al., 2017) with each container’s total polyp-days of survival as an offset, so containers that lost starved polyps, and thus had fewer polyp-days in which to reproduce, were compared on a per-polyp-day basis. Because this adjustment assumes a steady per-polyp rate over life, and unfed polyps declined before dying, it if anything understates the starved rate, so the reported gap between the fed and unfed rates is an upper bound. We compared the resulting per-polyp rates among schedules (R package emmeans 2.0.3, Lenth and Piaskowski, 2026); because a non-significant difference is not by itself evidence that the schedules were alike, we also tested whether their laceration rates fell within a pre-set 1.5-fold band of one another (Lakens, 2017). Model choice and fit were checked with information criteria and residual diagnostics (R package DHARMa 0.4.7, Hartig, 2024).

To test whether laceration tracked the feeding schedule, we modeled each cyclic schedule’s daily lacerate counts against the number of days since the last meal, again with a negative-binomial model that let each container have its own baseline. A rhythm shows up as a difference in laceration among the days of the cycle; we measured its strength with a likelihood-ratio test of how much this day-since-feeding term improved the model, and used the daily-fed animals, which never go more than a day without food, as a rhythm-free control. We also confirmed that the rhythm was not an artifact of treating successive daily counts within a container as independent, by refitting each model with a first-order autoregressive term for temporal correlation. Finally, we asked whether feeding history left a mark on the rhythm by testing whether an anemone’s first-trial schedule altered its second-trial cycle. Feeding history was tested in four ways altogether, on the total number of lacerates, on its interaction with the current schedule, and on each of the two cyclic rhythms, and we adjusted these four p-values together with the Holm procedure. No anemones died during the second trial, so its laceration totals needed no survival offset.

## 3. Results

Growth rose with feeding, but not without limit (Fig. 1). Anemones fed daily and every second day grew fastest and at similar rates (0.091 and 0.077 mm^2^ day^-1^; daily versus every second day, per-container growth rates, *t*(18) = 1.63*, p* = 0.122), so halving the number of meals from daily to every second day did not detectably slow growth. This similarity held for proportional as well as absolute growth and after adjusting for starting size. Growth fell off at the leaner schedules, with anemones fed every fourth day growing about half as fast (0.035 mm^2^ day^-1^) and unfed anemones shrinking. Over the 35 days, animals fed daily or every second day more than tripled in PDSA, those fed every fourth day doubled, and unfed survivors ended about a fifth smaller than they began. Because the schedules began at slightly different sizes (mean PDSA 1.07 to 1.33 mm^2^, unfed animals largest), we compared growth as the rate of area change rather than as final size.

**Figure 1.**
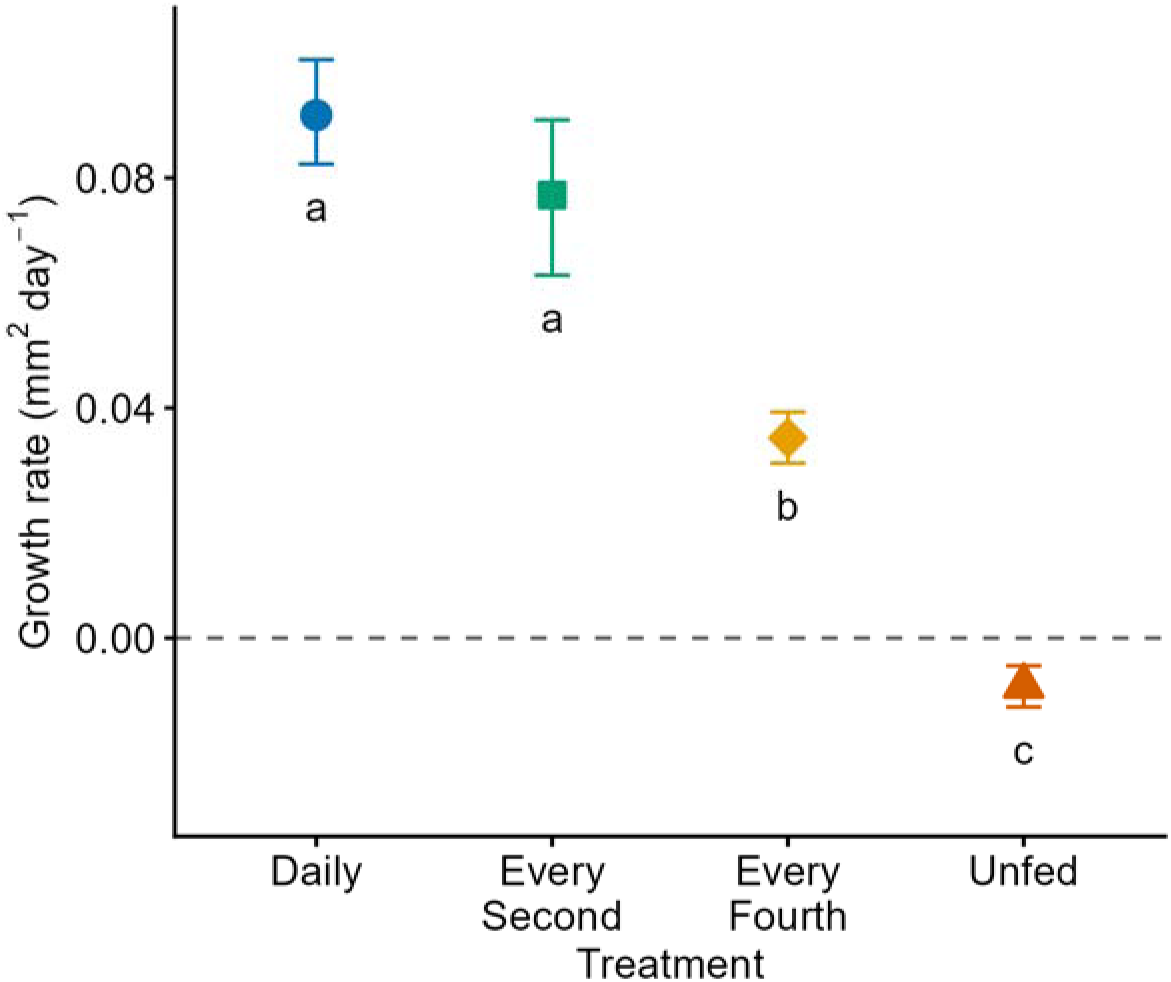
Growth rate (change in pedal disk surface area) of *Cylista elegans* under four feeding schedules in the first trial; points are the mean of per-container slopes with bootstrapped 95% confidence intervals. Shared letters below each point indicate schedules that did not statistically differ.

The number of lacerates depended on whether an anemone was fed rather than on how often (Fig. 2). The three fed schedules produced similar numbers of lacerates, about 19 to 23 per 100 polyp-days once we accounted for how long each polyp survived, with no detectable difference among them (*z* ≤ 1.72, *p* ≥ 0.317). An equivalence test placed two of the three pairwise rate ratios within a pre-set 1.5-fold margin; the exception was the daily versus every-second-day pair, whose confidence interval extended just past the margin, so equivalence was not established for that pair. Every fed rate exceeded the starved rate of 13 per 100 polyp-days (rate ratios 1.45 to 1.74, *z* ≥ 3.13, *p* ≤ 0.009).

**Figure 2.**
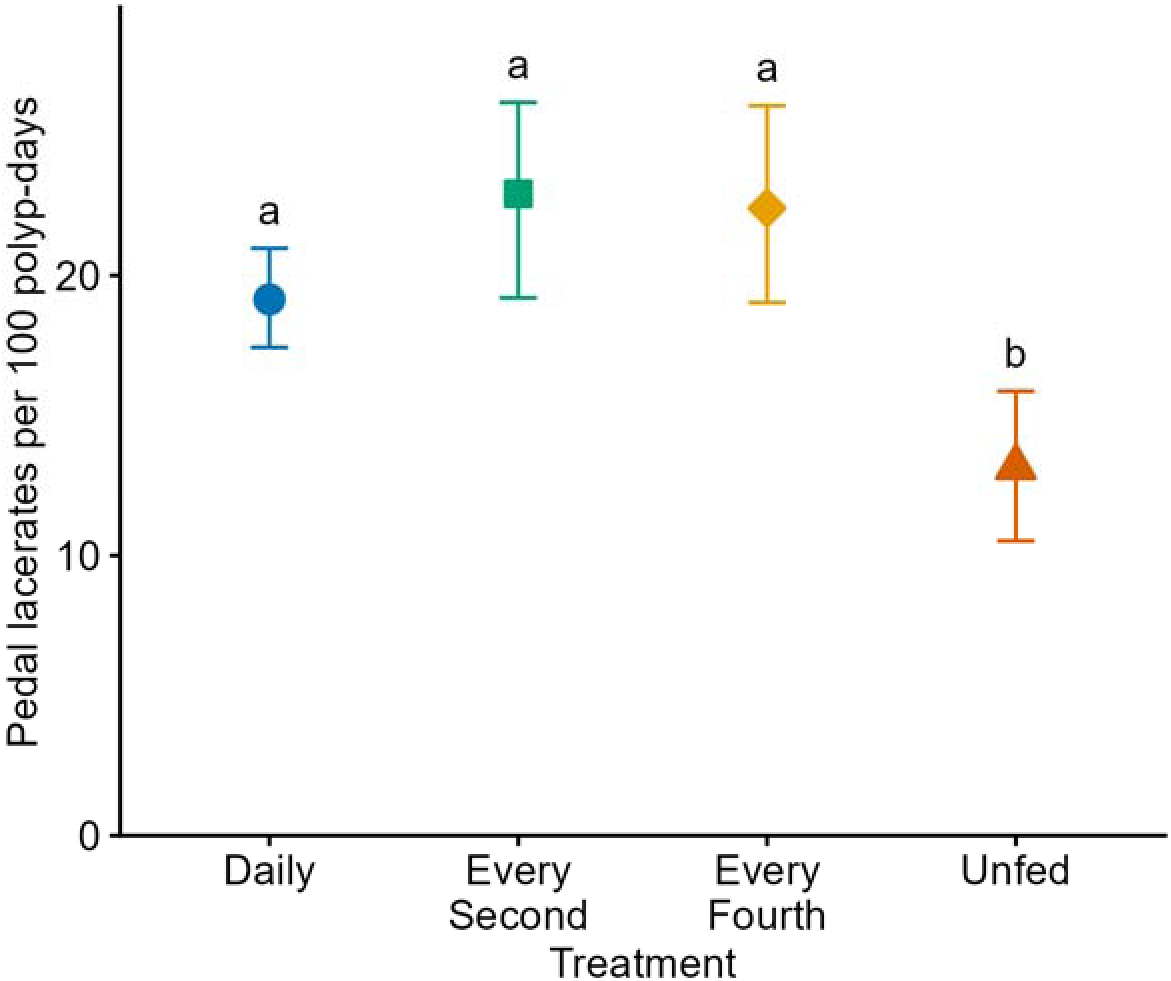
Pedal lacerates produced per 100 polyp-days under each feeding schedule in the first trial, with bootstrapped 95% confidence intervals. Shared letters above each point indicate schedules that did not statistically differ.

While how often an anemone was fed did not change how much it lacerated, the timing of feeding set the timing of laceration (Figs. 3, 4). Within each cycle, laceration fell to its lowest in the day after a meal and climbed as the interval since the last meal lengthened. Under every-fourth-day feeding the swing was wide. Production nearly stopped the day after feeding (0.04 lacerates polyp^-1^ day^-1^) and climbed almost tenfold to a peak on the third day after feeding (0.37 lacerates polyp^-1^ day^-1^). Under every-second-day feeding, where a two-day cycle allows only a single contrast, anemones lacerated less the day after a meal (0.17 lacerates polyp^-1^ day^-1^) than on the next feeding day itself, counted before its food (0.29 lacerates polyp^-1^ day^-1^). Daily-fed anemones, never more than a day from a meal, kept no such rhythm. The daily record makes the pattern clear (Fig. 4): production cycled on a two-day period under every-second-day feeding and a four-day period under every-fourth-day feeding, and was arrhythmic under daily feeding. Both cyclic schedules showed strong rhythms (every second day, χ^2^(1) = 15.9; every fourth day, χ^2^(3) = 125.1; both *p* < 0.001) and the daily schedule none (χ^2^(1) = 0.90, *p* = 0.344).Allowing for temporal autocorrelation within containers left these rhythms essentially unchanged (every second day, χ^2^(1) = 17.1; every fourth day, χ^2^(3) = 124.5; daily, *p* = 0.344).

**Figure 3.**
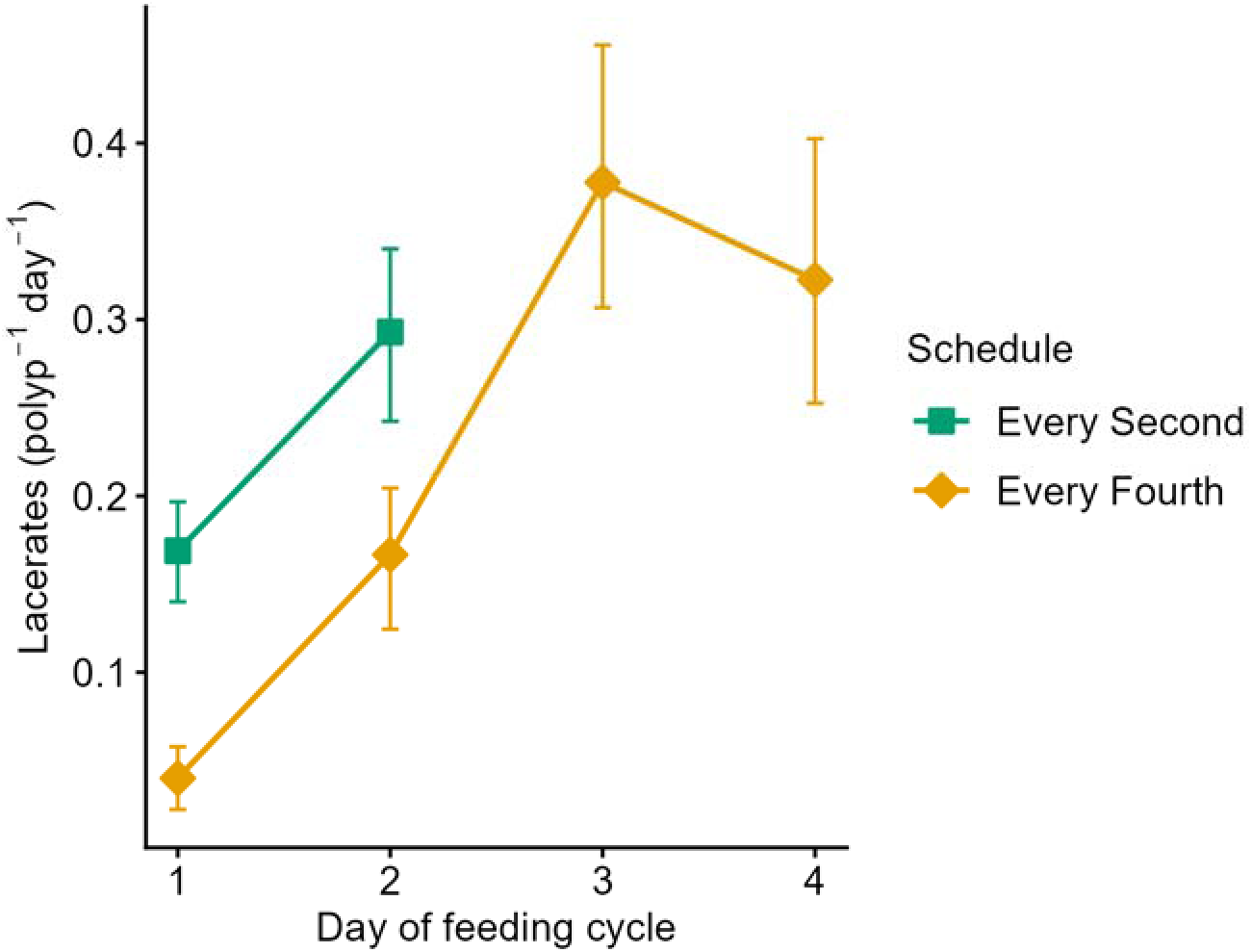
Pedal laceration by day of the feeding cycle, for the two cyclic schedules. Day 1 is the first 24 h after a meal; the last day of each cycle (day 2 for every-second-day feeding, day 4 for every-fourth-day feeding) is the count taken just before the next meal. Points are means with bootstrapped 95% confidence intervals.

**Figure 4.**
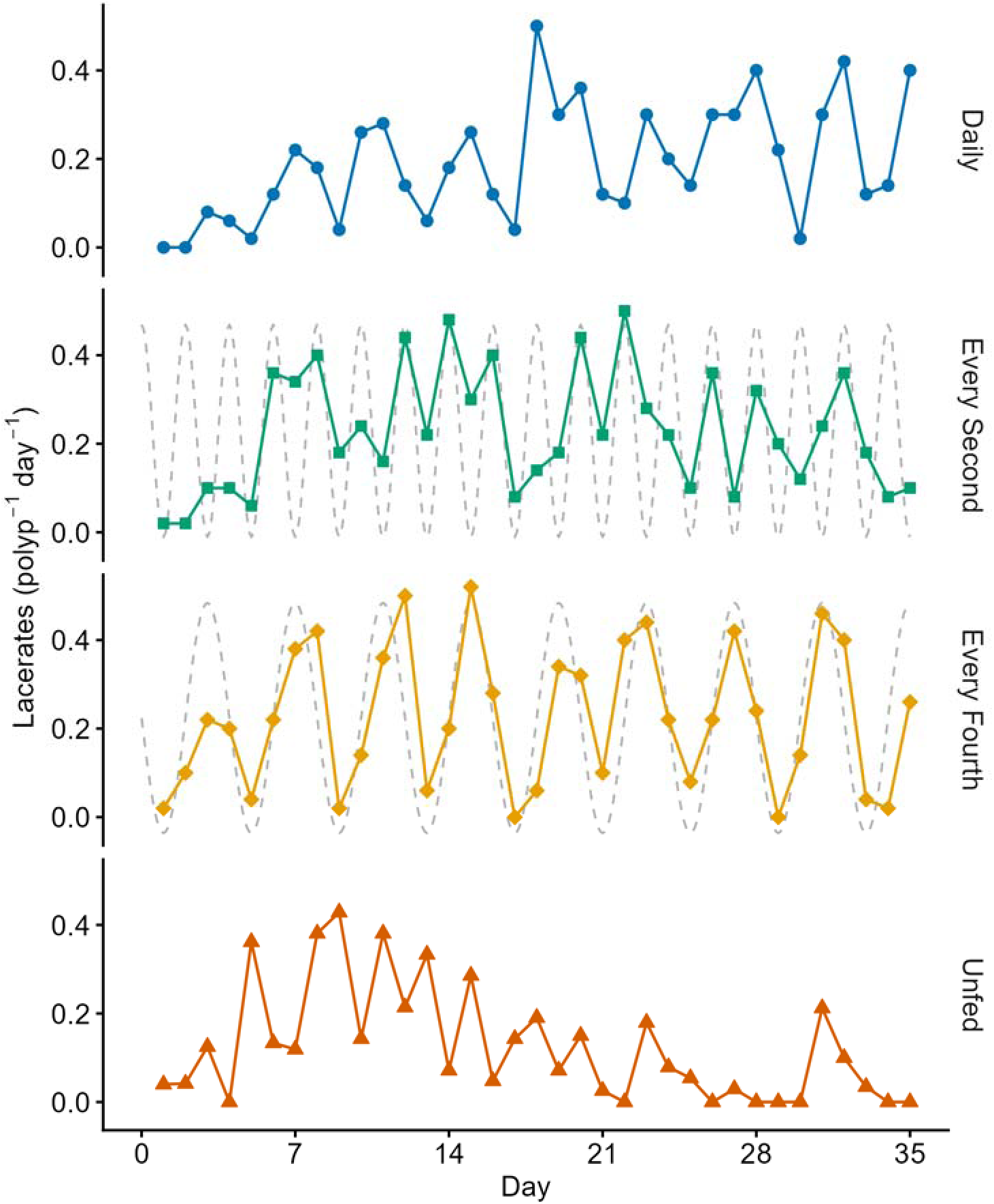
Daily pedal laceration through the 35 days of the first trial under each schedule. Dashed gray waves are a visual aid for tracing the two-day and four-day feeding cycle with feeding happening at the bottom of the wave.

Deaths occurred only under starvation (Fig. 5). Every anemone fed on any schedule survived the 35 days, whereas 21 of the 50 unfed anemones died.

**Figure 5.**
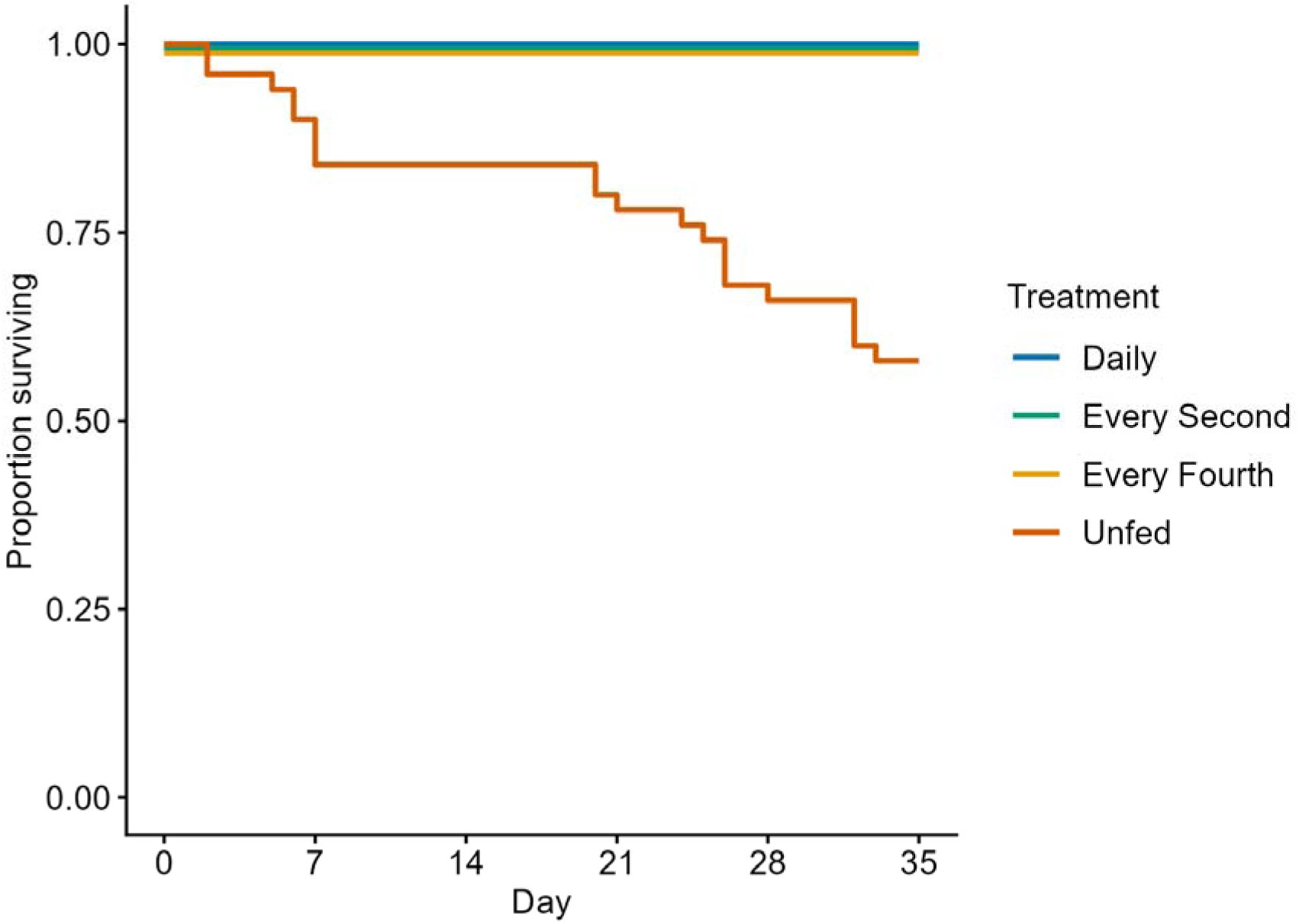
Kaplan-Meier survival under each feeding schedule in the first trial. All three fed schedules survived to day 35 and overlie one another at 1.0; their curves are offset slightly so all four are visible.

Reassigning anemones to new schedules for a second 35 days separated the effects of the current schedule from those of the previous one. Growth followed the new schedule, as in the first trial. Anemones moved to daily or every-second-day feeding grew, and those moved to leaner schedules shrank. Prior feeding appeared to affect growth as well, animals fed less in the first trial growing faster in the second (Fig. 6); this difference, however, reflected starting size rather than a lasting effect of feeding, since better-fed animals entered the second trial larger, and larger animals grew more slowly as they neared the growth ceiling (starting size and growth rate were inversely related, *r* = −0.48), and it vanished once starting size was included as a covariate (*F*(2,23) = 0.97, *p* = 0.395). Reproduction carried a clearer mark of the past. Animals fed daily or every second day in the first trial went on to lacerate more than those previously fed every fourth day, whatever their new schedule (main effect of prior feeding, χ^2^(2) = 13.43, *p* = 0.001). Whether this carryover also reshaped the response to the current schedule was less certain: laceration under every-fourth-day feeding was highest in animals earlier fed daily or every second day, but this interaction did not survive correction for the several second-trial history tests (interaction χ^2^(6) = 12.85, *p* = 0.045; Holm-adjusted *p* = 0.092).

**Figure 6.**
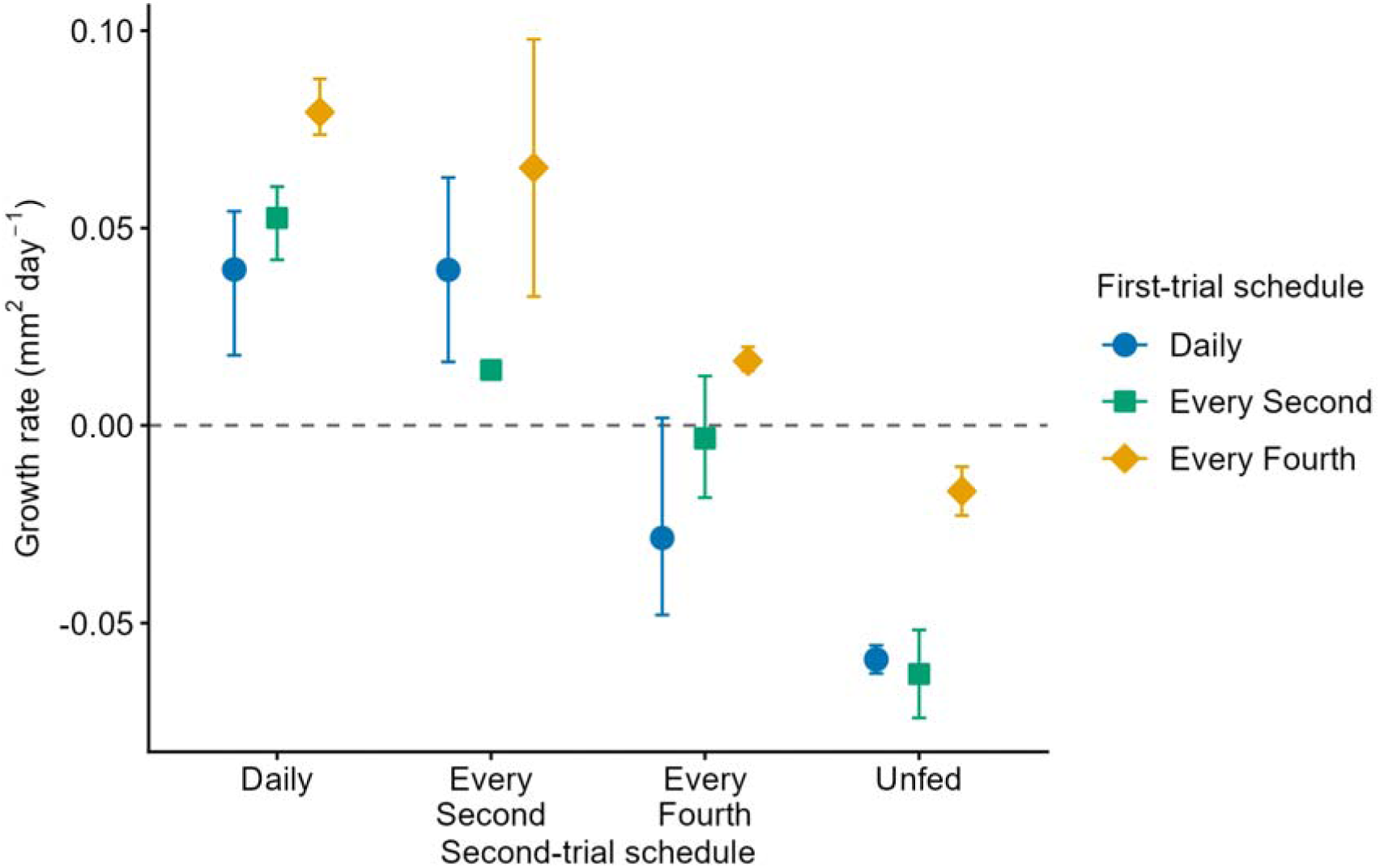
Second-trial growth rate against the second-trial feeding schedule, colored by the first-trial schedule; points are the mean of per-container slopes with bootstrapped 95% confidence intervals.

The rhythm’s period, unlike its magnitude, reset to the current schedule. Anemones moved to every-second-day feeding took up a two-day cycle and those moved to every-fourth-day feeding a four-day cycle, whatever schedule they had come from (every second day, χ^2^(1) = 59.3; every fourth day, χ^2^(3) = 112.1; both *p* < 0.001). Past feeding did not affect the every-second-day cycle (χ^2^(4) = 2.76, *p* = 0.599), and its apparent effect on the every-fourth-day cycle did not survive correction for the several second-trial history tests (χ^2^(8) = 16.95, *p* = 0.031, Holm-adjusted *p* = 0.092): an anemone lacerated on the period set by its current schedule, with no reliable trace of how it had been fed before.

## 4. Discussion

Feeding frequency shaped three aspects of asexual reproduction in *C. elegans*, and the three varied independently: the rate of growth, the number of lacerates, and the timing of laceration. The first two effects resemble those in other anemones, most directly *E*. *diaphana*. The third, a rhythm of laceration that matched the feeding schedule and reset whenever that schedule changed, is the central result of this study.

Growth rose with feeding frequency but saturated short of the most frequent schedule. Anemones fed daily grew no faster than those fed every second day, and each meal under daily feeding produced less growth than each meal under the less frequent schedules, so the most frequent feeding used food the least efficiently. A sea anemone has a blind gut, a single gastrovascular cavity that both takes in food and voids waste, so it processes each meal as a batch rather than continuously. A meal is digested over an interval approaching a day (Van-Praët, 1985; Zamer et al., 1987), so food arriving before the previous meal clears adds little. Food delivered more often than every second day yielded neither faster growth nor more lacerates. Individual growth rose with feeding in much the same way in *E. diaphana* (Clayton and Lasker, 1985), and the ceiling we observed indicates that these anemones were close to the fastest growth their bodies could sustain on a diet of brine shrimp, a ceiling set by the diet rather than by the size the animals could ultimately attain.

Total laceration, by contrast, depended only on whether an anemone was fed at all, not on how often. The same decoupling of total clonal output from the amount of food appears in the symbiotic anemone *E. diaphana*, where feeding rate changes the timing of laceration over the course of the experiment but not the number of lacerates produced (Clayton and Lasker, 1985; Hunter, 1984). The direction in which feeding drives reproduction, however, differs among species. In *E. diaphana* laceration is inversely related to feeding, rising under starvation and suppressed by feeding, and in *A. elegantissima* fission occurs mainly in the food-poor season and only in larger individuals in the field (Clayton, 1985; Sebens, 1980). The *A. elegantissima* pattern is not settled, though: a later factorial experiment found that fed anemones divided more often than unfed ones, with no effect of manipulating their symbionts (Tsuchida and Potts, 1994). *C. elegans* did the reverse, lacerating more overall when fed than when starved — though feeding suppressed laceration transiently in the day after each meal, and prolonged starvation eventually eroded reproduction along with growth — placing it nearer the feeding-stimulated fission of *D. lineata* and clone formation of the moon jelly *A. aurita* (Keen and Gong, 1989; Minasian, 1976). Whether symbiosis sets this direction remains doubtful: the one experiment to manipulate it found symbionts had no effect on fission in *A. elegantissima* (Tsuchida and Potts, 1994), and symbiotic and asymbiotic anemones differ in phylogeny and life history besides. What the contrast does establish is that the direction in which feeding drives asexual reproduction is itself labile, differing among the anemones in which it has been measured.

Whether an anemone was fed, not how often, is what drove it to lacerate, and its growth was the balance left over, the scope for growth that feeding provided minus the tissue shed as lacerates. Beyond every-second-day feeding, both that scope and the laceration it paid for had reached a ceiling, so additional meals changed neither. Where feeding did redirect output was between growing and multiplying. As the schedule thinned from every second to every fourth day, growth fell by half while total laceration held steady, so an anemone facing leaner food preserved its clonal output even as its own growth slowed. Asexual reproduction in anemones is often read in this light, as a way to maximize the surplus of energy intake over cost and, when food is scarce, to spread a fixed amount of tissue across many small individuals rather than one large one, raising the clone’s total feeding surface area (Keen and Gong, 1989; Sebens, 1979, 1987), though smaller bodies carry a higher mass-specific metabolic cost, which limits the energetic benefit of subdividing into ever-smaller individuals (Sebens, 1981, 1982). A clonal anemone can in principle enlarge itself or multiply itself (Bucklin, 1987; Francis, 1979), and which it does is shaped by its surroundings. In *D. lineata*, temperature drives the rate of fission and, through it, the body size an animal holds and how it splits its effort between growth and reproduction (Ryan, 2018; Ryan et al., 2019). In *C. elegans* the same balance shifted with food rather than temperature. Food acted on how fast the anemones grew rather than on how much they lacerated, so the balance moved toward reproduction as growth slowed, not as laceration rose. Because dividing keeps an anemone small, and larger anemones produce more gametes, the most frequently dividing individuals in *D. lineata* are also the smallest and least sexually fecund (Ryan et al., 2019); the growth-versus-multiplication trade-off we measured is likely one face of a broader division of resources among growth, clonal spread, and gametes, the sexual part of which we did not follow. Our anemones never reached the size at which *C. elegans* matures sexually, which in our field observations exceeds about a centimeter and which a diet of brine shrimp is unlikely to support. Converting limited and patchy food into more small, mobile clonemates that can disperse and feed independently, rather than into a single larger body, could be an adaptive response for an animal with a patchy, unpredictable food supply.

The timing of laceration followed the feeding schedule closely, lowest in the day after a meal and rising to a peak late in the cycle, so anemones fed every second or every fourth day lacerated on matching two- or four-day rhythms, while daily-fed animals, always digesting, kept no rhythm at all. Laceration thus concentrated in the non-feeding part of each cycle, the same timing Smith and Lenhoff (1976) found in *A. hyalinus* and unlike the day-to-day randomness of *E. diaphana* (Clayton, 1985). The suppression after feeding may reflect a partial conflict between laceration and digestion. Pedal laceration tears a fragment from the margin of the pedal disk, carrying away part of the endoderm and opening the gastrovascular cavity (Cary, 1911), though far less than longitudinal fission, which divides the cavity along its length. Laceration while a meal is being processed could interrupt digestion, so holding laceration until digestion is largely finished would avoid that conflict; the deeper disruption of fission would be even harder to reconcile with active digestion. In *Metridium senile*, the elevated metabolism that follows a meal subsides within about a day (Zamer et al., 1987), the interval over which laceration recovered here, so the suppression coincides with the period of active digestion (Van-Praët, 1985). That the every-fourth-day schedule produced a graded four-day rise rather than a simple alternation, with the minimum fixed on the day after feeding, ties the suppression to the timing of meals. Because laceration takes only a small fraction of an anemone’s energy budget (Hunter, 1984) and, unlike fission, leaves the parent able to keep feeding (Bucklin, 1987), any energetic cost would lie in processing the meal rather than in laceration itself. Both a mechanical conflict with active digestion and a temporary diversion of resources to the meal remain plausible.

When anemones were moved to a new schedule, the period of laceration changed to match it: an animal moved to every-second-day feeding took up a two-day cycle and one moved to every-fourth-day feeding a four-day cycle, regardless of the schedule it had come from. The timing of laceration thus followed current feeding rather than past feeding. Whether the rhythm is a direct response to how recently an anemone has eaten or an endogenous oscillator entrained by feeding cannot be settled here, because we did not test whether any cycle persists once feeding stops. Anemones do possess endogenous circadian rhythms of activity and metabolism that continue for days after the entraining cue is removed (Hendricks et al., 2012; Maas et al., 2016; Oren et al., 2015), yet in the anemone *Nematostella vectensis*, temperature cycles alone can drive rhythmic behavior, and setting light and temperature in conflict disrupts the endogenous clock itself (Berger and Tarrant, 2023); distinguishing an entrained clock from a direct response to feeding for laceration will require following the rhythm into constant conditions.

The total amount an anemone lacerated, by contrast, carried a mark of its past. The animals that had been fed most in the first trial went on to lacerate more whatever their new schedule, as though spending down a reserve built under the richer earlier regime. The tissue carried off in each lacerate is taken from stores in the pedal disk and replaced by feeding (White et al., 2026), and larger anemones, with more pedal-disk margin from which to shed, produce more lacerates (Bedgood et al., 2020), so a larger, better-provisioned anemone can keep producing clonemates for longer. Feeding history thus left the period of laceration untouched, resetting it to the current schedule, but shaped its magnitude, which followed the size and energy reserves the animal carried forward.

In *C. elegans*, then, asexual reproduction is bound closely to feeding: an anemone lacerates more when fed than when starved, holds laceration through the day after a meal, resumes it before the next, and lets growth, not laceration, track the food supply. This coupling places *C. elegans* among the clonal anemones whose asexual output rises with feeding, a group that includes the most successful of all anemone invaders, *D. lineata*. Because pedal laceration needs no mate, a single founder can start a population on its own. In practice, though, introduced *D. lineata*, now recorded as far south as Argentina (González Muñoz et al., 2023), usually carry several co-occurring genotypes, so their clonal reproduction contributes mainly by producing colonizing propagules rather than by letting one genotype fill a site (Ting and Geller, 2000). On that strength, together with a broad tolerance of temperature and salinity, the species has spread on ship hulls and oyster shells to become the most widely distributed sea anemone in the world (Hancock et al., 2017; Wood et al., 2022). The same clonality allows dispersal as well as population growth. Adults of *D. lineata* glide over surfaces and can detach and drift in currents away from poor conditions (Ryan et al., 2021), and *C. elegans* likewise releases from its substrate under thermal and food limitation stress (Wells, 2013; Wells and Harris, 2019). Yet *C. elegans*, which shares with *D. lineata* pedal laceration, a tolerance for the fouling communities of marinas, and this capacity for clonal spread, failed within a decade in its single western-Atlantic introduction (Wells and Harris, 2019).

Our results add a food axis to that comparison. If clonal output is tied as tightly to feeding in *C. elegans* as these experiments suggest, then the food supply of a candidate harbor, and not its temperature alone, may set whether the anemone can build the dense clonal population that establishment requires. Dock-dwelling anemones feed opportunistically on the local plankton, taking a broad range of prey (Wells et al., 2022), so the food available to an establishing population depends on the productivity of its particular harbor. Food is only one axis of a harbor’s suitability, alongside the physical setting, depth, light, and flow, that governs where temperate anemones live at all (Wells, 2019). A food-coupled invader like *C. elegans* should then establish only where food arrives reliably enough to keep starvation at bay, whereas *D. lineata*, whose fission responds to temperature as much as to food and so binds its clonal output less tightly to food supply, could persist on leaner or more erratic supplies. The contrast between the two, one failed and one now the most widespread sea anemone in the world, is well suited to test whether so tight a coupling of reproduction to food helps or hinders a clonal anemone in a new environment.

## CRediT authorship contribution statement

**Christopher D. Wells:** Conceptualization, Data curation, Formal analysis, Investigation, Methodology, Supervision, Validation, Visualization, Writing – original draft, Writing – review & editing. **Larry G. Harris:** Conceptualization, Methodology, Supervision, Writing – review & editing.

## Data availability

The cleaned datasets and the R and Quarto analysis and figure code that reproduce every result and figure in this article are openly available at Zenodo (https://doi.org/10.5281/zenodo.21342762) and on GitHub (https://github.com/christopherwells/cylista-feeding-frequency).

## Declaration of generative AI and AI-assisted technologies in the manuscript preparation process

During the preparation of this work the authors used Claude Opus 4.8 (Anthropic) to assess statistical analysis structure, organize content, and improve readability. After using this tool, the authors reviewed and edited the content and take full responsibility for the content of the published article.

## Acknowledgements

We thank the staff of the University of New Hampshire Department of Biology and Coastal Marine Laboratory for logistical support.

